# Insights into Yeast Response to Chemotherapeutic Agent through Time Series Genome-Scale Metabolic Models

**DOI:** 10.1101/2024.01.02.573982

**Authors:** Muhammed Erkan Karabekmez

**Affiliations:** Istanbul Medeniyet University, Department of Bioengineering

**Keywords:** Genome Scale Metabolic Models, Time Series Transcriptomics, Yeast, Doxorubicin

## Abstract

**Background:** Organism-specific genome-scale metabolic models (GSMMs) can unveil molecular mechanisms within cells and are commonly used in diverse applications, from synthetic biology, biotechnology, and systems biology to metabolic engineering. There are limited studies incorporating time-series transcriptomics in GSMM simulations. Yeast is an easy-to-manipulate model organism for tumor research;

**Methods:** Here, a novel approach (TS-GSMM) was proposed to integrate time-series transcriptomics with GSMMs to narrow down the feasible solution space of all possible flux distributions and attain time-series flux samples. The flux samples were clustered using machine learning techniques, and the clusters’ functional analysis was performed using reaction set enrichment analysis;

**Results:** A time series transcriptomics response of Yeast cells to a chemotherapeutic reagent – doxorubicin - was mapped onto a Yeast GSMM. Eleven flux clusters were obtained with our approach, and pathway dynamics were displayed. Induction of fluxes related to bicarbonate formation and transport, ergosterol and spermidine transport, and ATP production were captured;

**Conclusions:** Integrating time-series transcriptomics data with GSMMs is a promising approach to reveal pathway dynamics without any kinetic modeling and detects pathways that cannot be identified through transcriptomics-only analysis. The codes are available at https://github.com/karabekmez/TS-GSMM.

## 1. Introduction

Dynamics of metabolic pathways can be elucidated by different methods. Kinetic modeling is the most plausible approach at first sight but the excessive number of kinetic constants brought complications for this type of approaches. In order to reduce the complexity some efforts focus on simplifying genome scale metabolic models (GSMM) to enable targeted kinetic models [1]. GSMMs, firstly used to estimate flux distributions in steady state (FBA: Flux Balance Analysis; pFBA: Parsimonious FBA) [2–4] and to predict pseudo dynamic flux behaviors (dFBA: Dynamic FBA) [5–7]. Integrative analysis of transcriptomics and GSMMs is also common especially in higher organisms in order to make use of tissue or cell type specific GSMMs [8,9]. In some studies, conventional metabolic flux analysis is used by adding dynamic metabolomics [10,11] or dynamic transcriptomics [12] into the picture. Dynamic metabolomics can also integrated with GSMMs by manipulating constraints (uFBA) [13] or objective function (Metab_FBA) [14] considering each time point separately, however time-series metabolomic dataset are rare. The study by Hastings et al., builds time point specific GSMMs by using time-series transcriptomics towards E-Flux method [15] before using dynamic metabolomics in the objective function [14]. In another recent study, time point specific GSMMs were constructed by using GIMME approach [16] by integrating dynamic transcriptomics and then FBA was utilized to simulate growth characteristics [17]. However, GIMME approach yields almost the same network for each time step and flux variability should also be accounted. E-Flux sounds more plausible approach as it only manipulates flux bounds instead of eliminating reactions from the network. However, focusing on single flux distribution either by FBA or Metab_FBA might miss the flux variability within single time point. Additionally, in many cases cells grow sub-optimally, especially in stress conditions results of FBA should be handled cautiously and alternative objective functions should be employed [18]. Sampling the whole feasible solution space without searching any optimality is another common approach in GSMM related flux analysis [19]. Such Bayesian inference is promising if the solution space is narrow down by integrating omics data [20]. There are multiple algorithms for sampling the solution space; the artificially centered hit-and-run (ACHR) algorithm [21] and the coordinate hit-and-run with rounding (CHRR) algorithms [22] are sampling around optimal biomass production while Bayesian FBA (bFBA) [23] relaxes growth optimality to sample with a higher variance. The most important challenge in sampling is unreasonable run times for large GSMMs [24].

In this study, a pipeline called Time Series Genome Scale Metabolic Modeling (TS-GSMM) was proposed. Firstly, a modified version of E-Flux, fE-Flux [25], which guarantees feasibility of the resulting models, was employed to construct time step-specific constrained GSMMs from a Yeast GSMM by integrating a time series transcriptomic dataset collected after exposure to doxorubicin in a chemostat environment. ACHR approach was used to sample the flux cone for each time point, and the Wasserstein distance was used to create a distance matrix of time points for each reaction. The normalized distances of the resulting matrices were clustered using a k-medoid clustering approach. The obtained pseudodynamic flux clusters were functionally annotated through reaction set enrichment analysis.

## 2. Materials and Methods

A recent and moderate size GSMM for yeast, namely Yeast6 [26] was retrieved. Yeast6, v6.0.0 has 1882 reactions, 1454 metabolites and 901 genes out of which 881 metabolic genes’ expression values present in the transcriptomic dataset. The transcriptomic dataset consists of a time series response of yeast cell to a common therapeutic agent, doxorubicin in a chemostat [27]. The dataset can be reached from Array Express by accession number E-MTAB-10023.

The steps for TS-GSMM were conducted by using Cobra Toolbox (v.3.0) [28] in Matlab. Reaction boundaries in the Yeast6 model were adjusted by using Flux Variability Analysis (FVA) [29] letting 80% of the optimal objective function – biomass production possible, in order to be able to simulate sub-optimality under stress conditions. For each time point of the time-series transcriptomic dataset, reaction expressions were calculated by using GPR rules and metabolic gene expression values. *mapExpressionToReactions* function in the Cobra Toolbox was used for this purpose.

As proposed in E-Flux approach [15] adjusted reaction boundaries in the Yeast6 model was narrowed down proportional to the ratio of normalized expression values to the highest expression level in the same time point. However, these reaction boundary manipulations may lead to an infeasible GSMM. In order to prevent infeasibility, fE-Flux which is a modified version of E-Flux was implemented. fE-Flux skips reaction bound manipulations that leads to an infeasible model [25].

The artificially centered hit-and-run (ACHR) algorithm [21] was applied to span feasible solution space within time point specific GSMMs by using *sampleCbModel* function in the Cobra Toolbox. 2000 flux points were sampled for each reaction within each GSMM.

Pairwise distances of sampling distributions between time points were calculated by 1-Wasserstein distance [30] in Matlab. p-norm distance between generated distance matrices for each time series couple was assigned as flux profile distances. k-medoid clustering was used to get flux clusters out of it.

In order to decide the number of clusters for k-medoid clustering, Calinski-Harabasz clustering evaluation criterion (CH index) [31] and silhouette scores were calculated by using *evalclusters* function in Matlab R2022a [32]. In order to measure partitioning power of the number of clusters sum of squares of number of fluxes in each resulting cluster were also calculated.

Reaction set enrichment analysis were conducted by using KEGG [33] reaction networks for Yeast. Hypergeometric distribution was used to calculate p-values and Bonferroni correction was used to avoid multiple testing problem. 0.05 was used as threshold for corrected p-values.

Yeast metabolic pathways plots were retrieved from KEGG [33]. Box plots of flux samplings were obtained in Matlab R2022a.

## 3. Results

Yeast GSMM v6.0.0 has 1882 reactions, 1454 metabolites and 901 genes out of which 881 metabolic genes’ expression values present in the transcriptomic dataset. In conventional E-Flux applications the transcriptomics mapped GSMMs can be infeasible. In this study, all initially constructed time point specific GSMMs were found to be infeasible, i.e. no growth could be simulated. Therefore, we have used the fE-flux method. Initially FVA was applied by tolerating up to 80% of the optimal growth to find minimum and maximum flux boundaries (minFlux and maxFlux) for all fluxes in the reference Yeast6 GSMM. Then minFLux – maxFlux interval for each reaction in each time point was narrowed down proportional to the ratio of the corresponding reaction expression to the maximum reaction level for that time point. The reactions leading infeasibility were skipped.

It was observed that for 12 time point specific GSMMs that were constructed, upper bound adjustments of 8 to 10 reactions’ upper bound modifications led to infeasibility, out of which 8 reactions were the same for all time points. It means lowering those sensitive reactions’ upper bounds causes infeasibility. When we investigated the minFlux – maxFlux interval for those reactions and observed that typically all have a very narrow interval. The difference between the upper and lower bounds for these reactions are between 2,89E-5 and 1,26E-4, while 90% of the non-zero flux intervals are larger than this. ACHR sampling procedure was reduced time point specific GSMMs into models of identical reaction and metabolite sets of 965 reactions and 704 metabolites for ten time points. For the first two time points there were ten additional reactions which were excluded from the rest of the study. Pairwise distances of sampling distributions among 12 time points were calculated for 965 reactions by 1-Wasserstein distance for each time point. Unlike Kullback–Leibler divergence [34] or Kolmogorov–Smirnov statistic [35], Wasserstein distance give idea about true distance [36]. The distance matrix was clustered by k-medoids clustering approach assigning 11 as number of clusters which yields the highest number of clusters with seven or more fluxes as CH-index or silhouette scores were not helpful to decide the optimal number of clusters. The resulting numbers of fluxes in the clusters (Table 1) shows that majority of the fluxes resides in the same cluster (cluster 1) where there was no characteristic time series trend.

**Table 1.** The Numbers of fluxes in each cluster found by k-medoid approach.

| Cluster ID | 1 | 2 | 3 | 4 | 5 | 6 | 7 | 8 | 9 | 10 | 11 |
| --- | --- | --- | --- | --- | --- | --- | --- | --- | --- | --- | --- |
| Number of Fluxes in the cluster | 884 | 16 | 6 | 8 | 12 | 1 | 3 | 16 | 9 | 1 | 9 |

KEGG ID’s corresponding to reactions in the GSMM were already provided in Yeast6 but not all reactions have matches in KEGG like transport and exchange reactions. The clusters in which less than 3 KEGG ID’s are assigned were not used for reaction set enrichment analysis. Cluster 1, 2, 4 and 9 have more than 3 reactions with KEGG ID. Nucleotide and amino acid metabolism related pathways were found to be enriched in the cluster 1. The second cluster was found to be enriched in pyruvate metabolism and galactose metabolism is linked to the cluster 4. The pathway for biosynthesis of aromatic amino acids, phenylalanine tyrosine and tryptophan, was observed to be significantly associated with both cluster 1 and cluster 9 (Table 2).

**Table 2.** The Numbers of fluxes in each cluster found by k-medoid approach.

| CLUSTER 1 |  |  |  |
| --- | --- | --- | --- |
| KEGG Pathway | Corrected<br>p-values | Number<br>of<br>Reactions | Number<br>of<br>Matches |
| Pyrimidine metabolism | 0,00E+00 | 100 | 32 |
| Purine metabolism | 0,00E+00 | 154 | 45 |
| Cysteine and methionine metabolism | 3,98E-10 | 100 | 22 |
| Alanine aspartate and glutamate metabolism | 3,29E-09 | 47 | 15 |
| Valine leucine and isoleucine biosynthesis | 2,36E-08 | 25 | 11 |
| Pyruvate metabolism | 4,53E-08 | 83 | 18 |
| Phenylalanine tyrosine and tryptophan biosynthesis | 5,68E-08 | 48 | 14 |
| One carbon pool by folate | 1,13E-06 | 34 | 11 |
| Glycine serine and threonine metabolism | 3,95E-06 | 75 | 15 |
| Carbon fixation in photosynthetic organisms | 1,65E-04 | 26 | 8 |
| Pantothenate and CoA biosynthesis | 2,04E-04 | 35 | 9 |
| Carbon fixation pathways in prokaryotes | 2,87E-04 | 56 | 11 |
| Citrate cycle (TCA cycle) | 3,22E-03 | 28 | 7 |
| Lysine biosynthesis | 4,30E-03 | 39 | 8 |
| Arginine biosynthesis | 6,50E-03 | 31 | 7 |
| Glycolysis / Gluconeogenesis | 1,28E-02 | 57 | 9 |
| Glycerophospholipid metabolism | 3,53E-02 | 93 | 11 |
| Histidine metabolism | 3,97E-02 | 53 | 8 |
| Steroid biosynthesis | 4,43E-02 | 67 | 9 |
| Fatty acid biosynthesis | 4,94E-02 | 68 | 9 |
| CLUSTER 2 |  |  |  |
| Carbon fixation in photosynthetic organisms | 6,21E-04 | 26 | 2 |
| Pyruvate metabolism | 6,44E-03 | 83 | 2 |
| CLUSTER 4 |  |  |  |
| Galactose metabolism | 2,25E-06 | 58 | 3 |
| Amino sugar and nucleotide sugar metabolism | 3,86E-05 | 148 | 3 |
| CLUSTER 9 |  |  |  |
| Phenylalanine tyrosine and tryptophan biosynthesis | 2,53E-04 | 48 | 2 |

Energy metabolism related fluxes are enriched in the cluster 2. Transport fluxes of malate-aspartate NADH shuttle were augmented in long term response. Ethanol processing and transport in mitochondria was also observed to be enhanced in a correlated trend (Figure 1).

**Figure 1.**
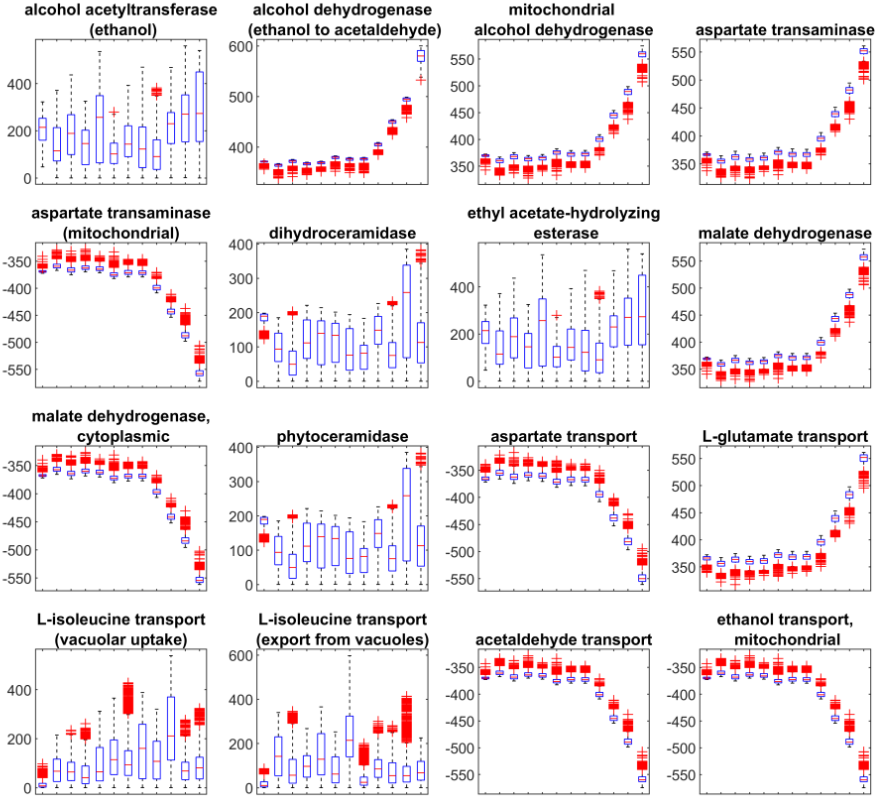
Flux sampling profiles in cluster 2. Each box plot represents flux sampling distribution at a time point.

All reactions in cluster 3 are related to bicarbonate formation (in cytoplasm and nucleus) and transport of carbon dioxide and water between nucleus and cytoplasm related to the bicarbonate formation reactions. Flux profiles shows that bicarbonate formation is shifted from nucleus to cytoplasm and there are two consecutive shifts in the reaction directions. After the changes in the reaction direction the initial state is reattained in the long term (Figure S1.).

There are 4 enzymatic reactions and 4 transport reactions in cluster 4. The enzymatic reactions are related to galactose metabolism (Table 2) while the rest are L-leucine and fatty acid transport associated transport reactions. Initial (before the pulse) flux distributions of these transport reactions are closer to zero and they tend to increase after the introduction of the chemotherapeutic (Figure S2.).

Ergosterol, spermidine and glycerol transport related fluxes reside in cluster 5. All these reactions seem to be induced upon introduction of doxorubicin into the chemostat environment (Figure S3.). Similar to cluster 3 and cluster 5, cluster 8 and cluster 11 were also found to be occupied dominantly by transport reactions. Increase in fluxes of AMP/ATP transporter, transport of ATP and some amino acids were indicated in the cluster 8 (Figure S4.). ATP synthase, ADP/ATP transporter, water diffusion and phosphate trans-port fluxes were observed to be induced in long term with a trend of flux distributions with much lower variances in the cluster 11. ATP synthase flux drops immediately after the pulse but increases in long term (Figure 2).

**Figure 2.**
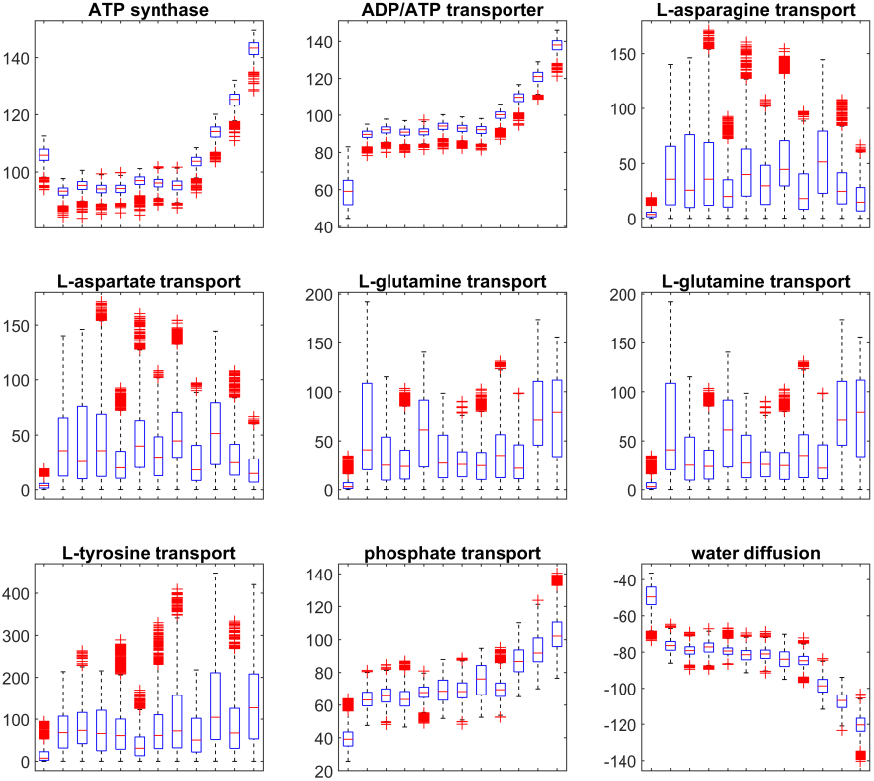
Flux sampling profiles in cluster 11. Each box plot represents flux sampling distribution at a time point.

Growth and glucose transport profiles are quite similar with a small initial increase and subsequent relatively higher decrease. Note that the difference between the top and bottom points is quite small though significant (Figure 3).

**Figure 3.**
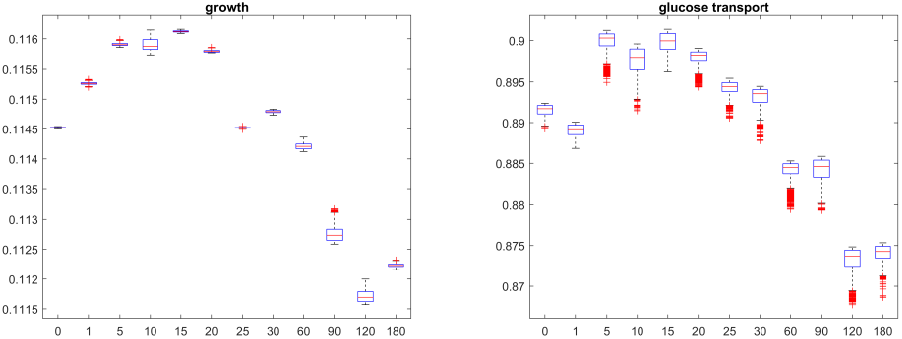
Flux sampling profiles of growth reaction (left) and glucose transport (right). x-axis is non-uniform time points in minutes.

When glycolysis pathway investigated in detail, it was observed that fluxes of four consecutive reactions from cluster 1; phosphoglycerate kinase, phosphoglycerate mutase, enolase and pyruvate kinase were sampled. Mean of flux values of phosphoglycerate kinase tend to decrease and that of pyruvate kinase tend to increase. The two reactions in between show no significant flux sampling distribution. It is interesting that reaction expression profiles give no clue for flux sampling profiles (Figure 4).

**Figure 4.**
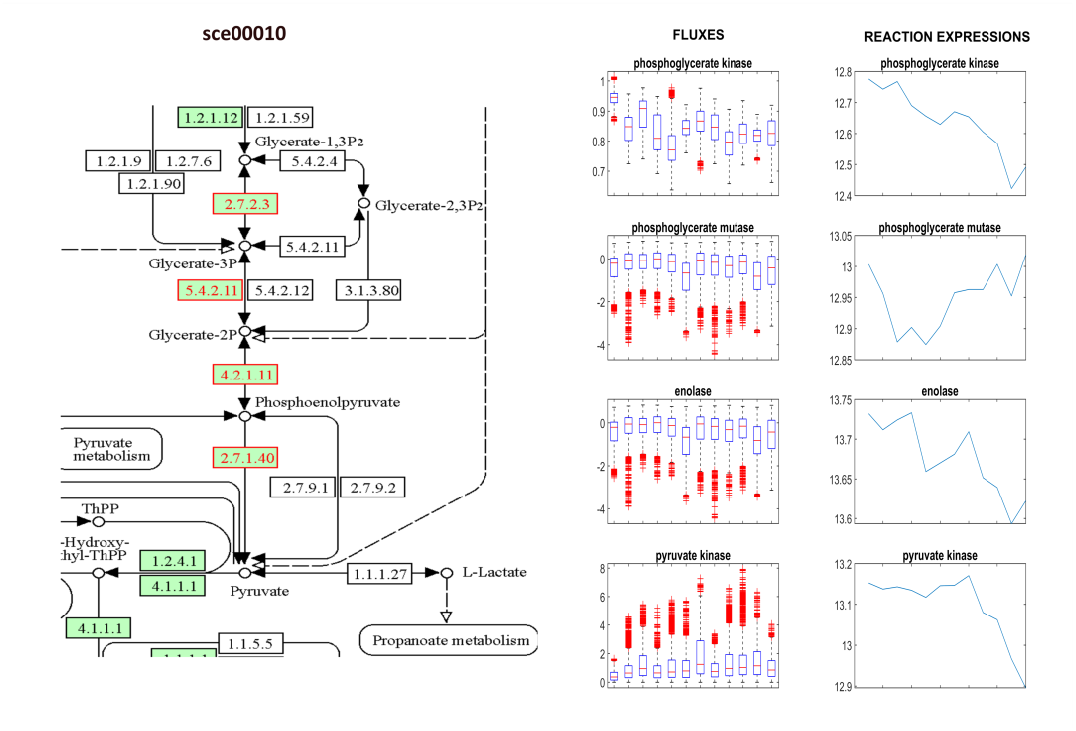
Flux sampling profiles of reactions glycolysis pathway. (sce00010: partial KEGG glycolysis pathway for yeast; FLUXES: flux sampling profiles for the reactions highlighted in red in the KEGG network with the same order. Each box plot represents flux sampling distribution at a time point; REACTION EXPRESSIONS: Reaction expressions obtained by mapping gene expressions to the GSMM by GPR rules)

## 4. Discussion

Malate-aspartate NADH shuttle (MAS) in yeast was reported to be one of the important mediators of extending life span [37]. It was also shown that aerobic glycolysis, which is one of the characteristic of tumor cells, can be limited by inducing MAS [38]. It is displayed in this study that MAS related transport fluxes were upregulated (Figure 1) upon exposure to doxorubicin. Interestingly this could help to build new hypotheses for lightening the mechanisms of anti-tumor activity of doxorubicin. Insufficiency of MAS could be related to heart failure [39] which is the most common side effect of doxorubicin. It is already reported that drugs inhibiting MAS can prevent cardiotoxicity of doxorubicin [40]. ATP production is also an important issue in cardio toxic side effect of doxorubicin. In human heart it is known that doxorubicin reduces ATP production efficiency [41]. Short term response of yeast cells might be simulating this phenomenon (Figure 2). In long term, yeast cells activate MAS and increases ATP production and related transport reactions.

In cluster 3, flux sampling profiles points a shift in directions of bicarbonate formation reactions resulting in down regulation of nucleolar bicarbonate formation and a reverse effect in the cytoplasm (Figure S1.). Bicarbonate formation has important roles in human diseases through its role in pH buffering [42]. It is also suggested that tumor cells may induce bicarbonate transport to regulate pH and resist to chemotherapeutics [43]. Bicarbonate’s possible role in cardiotoxicity of doxorubicin was also discussed in the literature [44].

Spermidine is claimed to induce autophagy of organelles in order to increase lifespan of yeast [45]. It can be hypothesized that doxorubicin leads to damages inside cells through formation of reactive oxygen species and yeast cells induce spermidine flux in response to this stress. Ergosterol plays a similar role in yeast with cholesterol’s role in human; it’s essential for membrane formation and maintenance. It is known that Ergosterol transport is enhanced in tumor cells in order to promote growth. Here, in this context a possible explanation is that – similar to cholesterol – yeast cells might be trying to reduce cellular doxorubicin concentration through ergosterol [46,47]. In this study, the flux changes detected in the cluster 5 captures induction in reactions related to ergosterol and spermidine transport (Figure S3.).

GSMM provides a template to reflect flux dynamics without involving reaction kinetics by integrating time series transcriptomics datasets. The proposed method captures pathways that cannot unveiled by analyzing solely transcriptomic data [27,48]. Future studies on time series proteomics will improve the performance of the proposed method.

## Supplementary Materials

Figure S1: Flux sampling profiles in cluster 3. Each box plot represents flux sampling distribution at a time point; Figure S2: Flux sampling profiles in cluster 4. Each box plot represents flux sampling distribution at a time point; Figure S3: Flux sampling profiles in cluster 5. Each box plot represents flux sampling distribution at a time point; Figure S4: Flux sampling profiles in cluster 8. Each box plot represents flux sampling distribution at a time point; Figure S5: Flux sampling profiles in cluster 9. Each box plot represents flux sampling distribution at a time point.

## Data Availability Statement

The codes for TS-GSMM and simulations were deposited at https://github.com/karabekmez/TS-GSMM.

## Acknowledgments

The author kindly express his gratitude to Prof. Dr. Mehmet Güray Güler, Furkan Cantürk and Merve Yarıcı for helpful discussions on technical details.

